# Evidence for enhancer noncoding RNAs (enhancer-ncRNAs) with gene regulatory functions relevant to neurodevelopmental disorders

**DOI:** 10.1101/2020.05.16.087395

**Authors:** Yazdan Asgari, Julian I.T. Heng, Nigel Lovell, Alistair R. R. Forrest, Hamid Alinejad-Rokny

**Author notes:** Correspondence should be addressed to H.A.R.

## Abstract

Noncoding RNAs (ncRNAs) comprise a significant proportion of the mammalian genome, but their biological significance in neurodevelopment disorders is poorly understood. In this study, we identified 908 brain-enriched noncoding RNAs comprising at least one nervous system-related eQTL polymorphism that is associated with protein coding genes and also overlap with chromatin states characterised as enhancers. We referred to such noncoding RNAs with putative enhancer activity as brain ‘enhancer-ncRNAs’. By integrating GWAS SNPs and Copy Number Variation (CNV) data from neurodevelopment disorders, we found that 265 enhancer-ncRNAs were either mutated (CNV deletion or duplication) or contain at least one GWAS SNPs in the context of such conditions. Of these, the eQTL-associated gene for 82 enhancer-ncRNAs did not overlap with either GWAS SNPs or CNVs suggesting in such contexts that mutations to neurodevelopment gene enhancers disrupt ncRNA interaction. Taken together, we identified 49 novel NDD-associated ncRNAs that influence genomic enhancers during neurodevelopment, suggesting enhancer mutations may be relevant to the functions for such ncRNAs in neurodevelopmental disorders.

## Introduction

Neurodevelopmental disorders (NDDs) represent a group of early onset disorders that arise as a result of defective nervous system development during prenatal and early postnatal stages. In NDDs which are associated with abnormal brain development, individuals may present with severe difficulties in emotional regulation, speech and language, autistic behavior, abnormalities of the endocrine system, attention deficit hyperactivity disorder, cognitive impairment, psychosis, and schizophrenia [1]. It is known that genetic, epigenetic, and environmental insults can disrupt the development of the central nervous system, including the birth of neural cells, their positioning and connectivity as functional units within brain circuitry, as well as their capacity for signal propagation [2]–[4]. As a corollary, NDDs arise as a consequence of defective neurogenesis, aberrant cell migration essential to neuronal positioning during circuit formation, as well as from dysfunctional synaptogenesis and aberrant synaptic pruning [5]. Despite significant advances in clinical insights into NDDs, the molecular mechanisms which underlie these often lifelong neurological conditions remains poorly characterised [1].

Our understanding of the genomic regulation of mammalian development, homeostasis and disease has been facilitated by significant technological advances in sequencing. As an example, since the discovery of noncoding RNAs (ncRNAs) [6], their diversity of subtypes, and their myriad roles in homeostasis and disease are becoming better understood. At present, ncRNAs can be classified into multiple groups, including transfer RNAs (tRNAs), ribosomal RNAs (rRNAs), microRNAs (miRNAs), Piwi-interacting RNAs (piRNAs), small interfering RNAs (siRNAs), small nucleolar RNAs (snoRNAs), small nuclear RNAs (snRNAs), extracellular RNAs (exRNAs), small cajal body-specific RNAs (scaRNAs), and long ncRNAs (lncRNAs) [7]. Of note, recent evidence is indicative of significant roles for ncRNAs in brain function and behavior. As a corollary, ncRNAs are implicated in the etiology of conditions including bipolar disorder, schizophrenia, autism, and language impairment [8][9]. Anecdotally, the functions for several ncRNAs species including H19, *KCNQ1OT1, SNORD115*, and *SNORD116*, all of which have been demonstrated to be relevant to neurodevelopment, are associated with NDDs [10][11]. Moreover, several large-scale studies have been characterised ncRNA functions in human neuropathological phenotypes [12][13][14], [15], yet the regulatory actions for ncRNAs in NDDs remains unclear.

In this study, we have investigated a curated set of noncoding RNAs for gene regulatory functions in neurodevelopment and NDDs, using an integrative bioinformatics pipeline. By integrating genomic and epigenetic datasets on these ncRNA species, analysing their ENCODE regulatory features, FANTOM5 tissue-specific enhancer traits, FANTOM5 expression profiles, nervous system-related expression quantitative trait loci (eQTLs) mapping, neurodevelopment-related GWAS single nucleotide polymorphisms (SNPs), and Copy Number Variation (CNV); we have identified a set of noncoding RNAs with evidence of gene regulatory functions in neurodevelopment and disease.

## Results

An overview of the pipeline used in this study is provided as **Figure 1**. As shown, we utilised the resource FANTOMCAT [16] to identify noncoding RNAs that are significantly enriched in brain tissue. Next, we annotated brain-enriched ncRNAs with ENCODE predicted chromatin states/histone modifications to identify promoters and enhancers for brain-enriched ncRNAs. These ncRNAs within enhancer loci are then investigated for their association with eQTLs to identify their linkage to a coding gene. Based on these criteria, we refer to these as neuronal ‘enhancer-ncRNAs’. Using data on CNVs and GWAS SNPs related to neurodevelopmental conditions, we then cross-referenced these datasets to identify enhancer ncRNAs that are mutated in individuals with NDDs, then focus on those enhancer ncRNAs in which an NDD mutation or CNV disrupts their association with a coding gene. Therefore, through these selection criteria, we identify a set of ncRNAs with putative gene regulatory roles relevant to NDDs.

**Figure 1.**
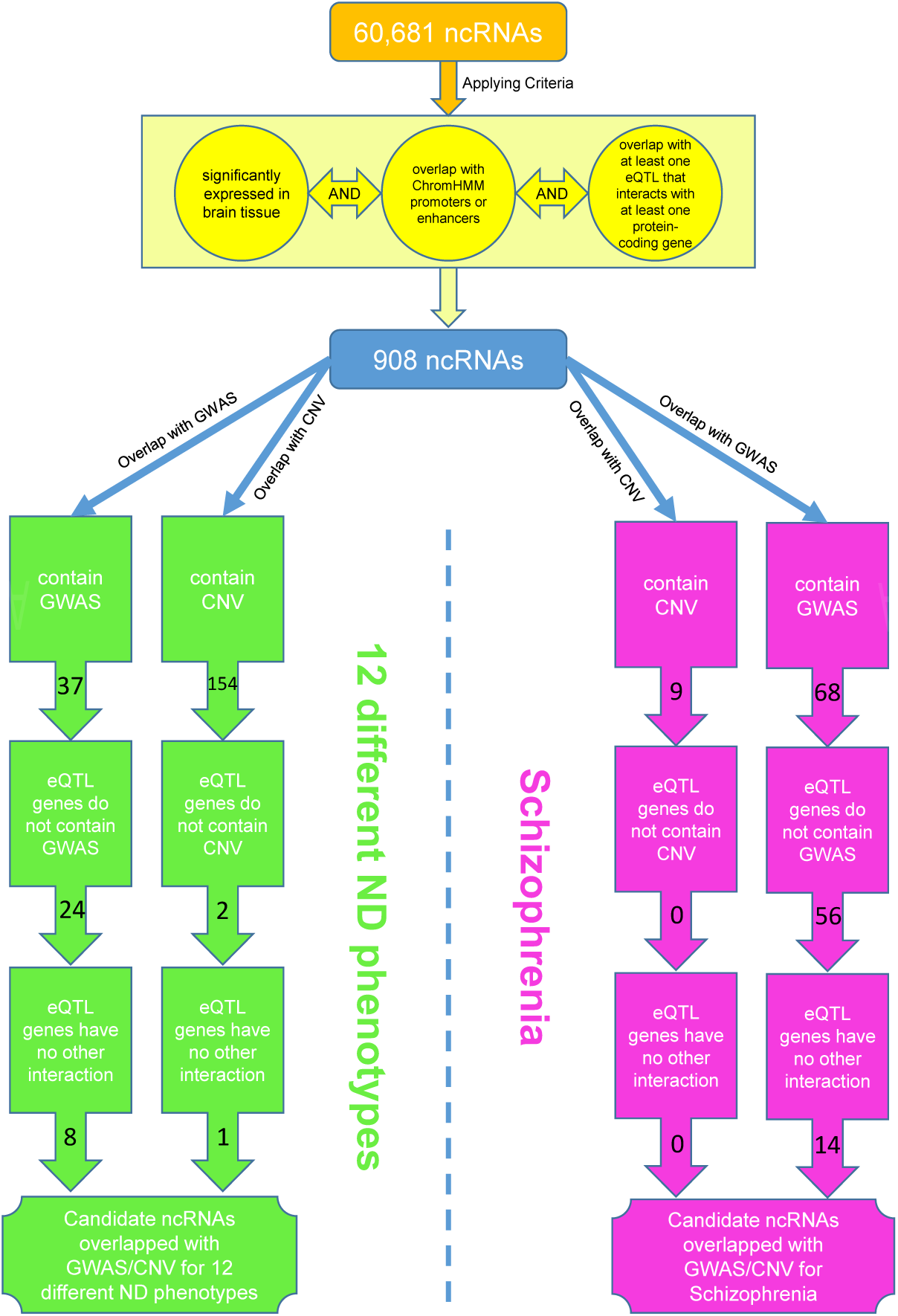
An overview of the integrative bioinformatics pipeline used in this study. Out of 60,681 noncoding genes in the list, we have started with selection of the ones with significant expression for brain tissue, overlapping with ENCODE predicted chromatin states presented in brain defined as ‘enhancer’ or ‘promoter’, and containing brain related eQTL polymorphism(s) that interacts with at least one protein-coding gene. Based on the above criteria, 908 noncoding RNAs annotated as putative enhancers in brain, focusing on to identify enhancer ncRNAs with possible regulatory function in neurodevelopmental disorders. We then explored the number of candidate ncRNA enhancers that contain NDD associated GWAS or overlap with NDD related CNVs whereas their eQTL genes do not contain GWAS/CNV. As a result, we identified 82 candidate ncRNA enhancers with potential regulatory function in neurodevelopmental disorders.

### Noncoding RNAs with histone mark and ChromHMM signals

We curated a set of noncoding RNAs comprising 60,681 noncoding genes including 31,343 long noncoding RNAs, from FANTOMCAT (**Figure 2a**). Brain-related ncRNAs were identified based on their mapping to ChromHMM-predicted promoter or enhancer loci. To do this, we selected genes corresponding to brain–specific promoter or enhancer regions (*see Methods for details*). Of 22,698 coding genes, 6,636 (29.24%) overlapped with promoters, 1,095 (4.82%) overlapped with enhancers, and 7,731 (34.06%) overlapped with either promoters or enhancers (**Figure 2b** and **Supplementary table S1**). Furthermore, of 60,681 noncoding genes, 5,736 of them (9.45%) overlapped with promoters, 2,945 (4.85%) overlapped with enhancers and 8,681 (14.31%) overlapped with either promoters or enhancers (**Figure 2c** and **Supplementary table S1**). In subsequent analyses, we only analyse noncoding genes that overlapped with either promoters or enhancers.

**Figure 2.**
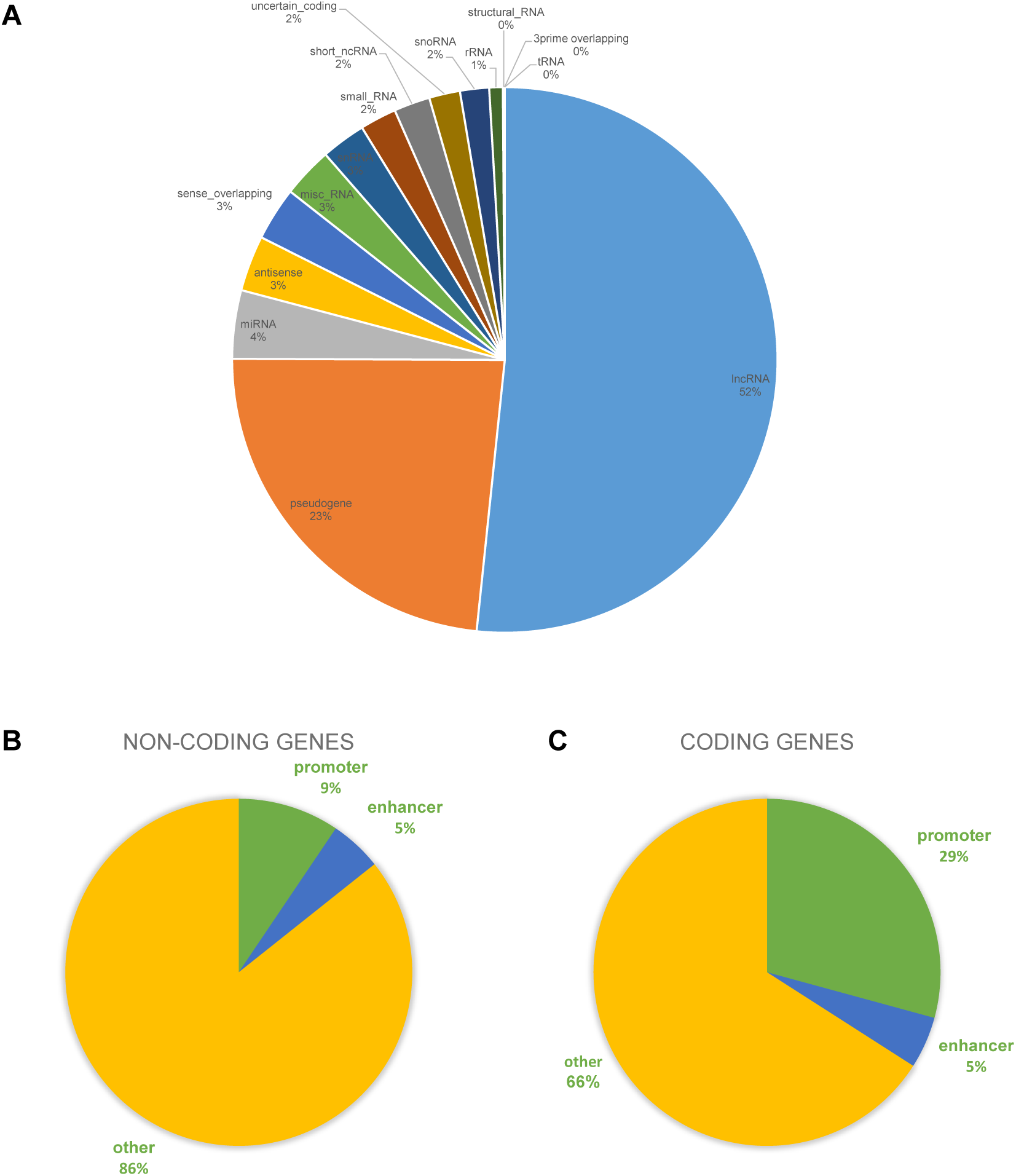
**A) Distribution of noncoding genes available in the gene list of this study.** Out of 84,065 genes available in the final list of the study, 60,681 are noncoding RNA genes. The figure shows fraction of each noncoding types through all 60,681 noncoding genes. The most fraction of ncRNAs are lncRNAs and pseudogenes while tRNAs, structural RNAs, and 3prime overlapping RNAs are the least ones. **B) Pie chart derived from ChromHMM data overlapped with ncRNAs**. Out of 60,681 noncoding genes, 9.45% overlapped with promoters, 4.85% overlapped with enhancers, and 14.31% overlapped with either promoters or enhancers. **C) Pie chart derived from ChromHMM data overlapped with coding genes**. Out of 22,698 coding genes, 29.24% overlapped with promoters, 4.82% overlapped with enhancers, and 34.06% overlapped with either promoters or enhancers.

### eQTL polymorphism suggests enhancer activity for ncRNAs

eQTLs are genomic loci that affect expression levels of genes in one or multiple tissues [17], [18], [19]. To identify noncoding RNAs that contain at least one brain-related eQTL polymorphism, we utilised data on eQTL pairs relevant to the nervous system from the Genotype-Tissue Expression (GTEx) Project [19].

When we performed this analysis with 60,681 noncoding genes, we found 17,743 (29.24%) that contained at least one eQTL polymorphism related to nervous system tissues, including 3,196 in Amygdala, 5,621 in Anterior Cingulate Cortex, 7,612 in Caudate Basal Ganglia, 8,682 in Cerebellar Hemisphere, 10,900 eQTL pairs in Cerebellum, 8,242 eQTL pairs in Cerebral Cortex, 6,476 eQTL pairs in Frontal Cortex, 4,462 eQTL pairs in Hippocampus, 4,503 eQTL pairs in Hypothalamus, 6,818 eQTL pairs in Nucleus Accumbens Basal Ganglia, 5,547 eQTL pairs in Putamen Basal Ganglia, 3,553 eQTL pairs in Spinal Cord Cervical, and 2,586 eQTL pairs in Substantia Nigra (**Supplementary table S2**).

We performed hierarchical clustering of 17,743 noncoding RNA genes that contained at least one nervous system related eQTL polymorphism based on their expression profiles. Through this approach, we found that a substantial proportion of genes are highly expressed in brain tissue, with smaller clusters of genes that are highly expressed in blood and T-cells, as well as granulocytes (**Figure 3**). This clustering also reveals a smaller, but nonetheless significant clustering of ncRNA genes highly expressed in testis, and within the male reproductive system (**Figure 3**, in green). Next, we investigated the distribution of eQTL polymorphisms across noncoding genes based on their ncRNA sub-classification. As shown in **Table 1**, across 13 different eQTL brain sub-regions, the majority of nervous system related eQTL polymorphisms are relevant to 17,743 lncRNAs (64.81%), followed by pseudogenes 21.93%, miRNAs 0.31%, antisense ncRNAs 4.68%, sense overlapping ncRNAs 2.09%, misc_RNAs 0.58%, snRNAs 0.25%, snoRNAs 0.24%, rRNAs 0.11%, 3prime overlapping ncRNAs 0.03%, but not tRNAs (0%). In addition, ncRNAs which do not sub-classify to these categories constitute a small proportion (4.96%). Thus, brain eQTL associated genes, on average, interact with 5.87 coding- and 3.18 non-coding genes that contain at least one brain related eQTL polymorphism (**Supplementary table S3**). This suggests that interactions between coding genes are preferred in nervous tissue.

**Table 1:**
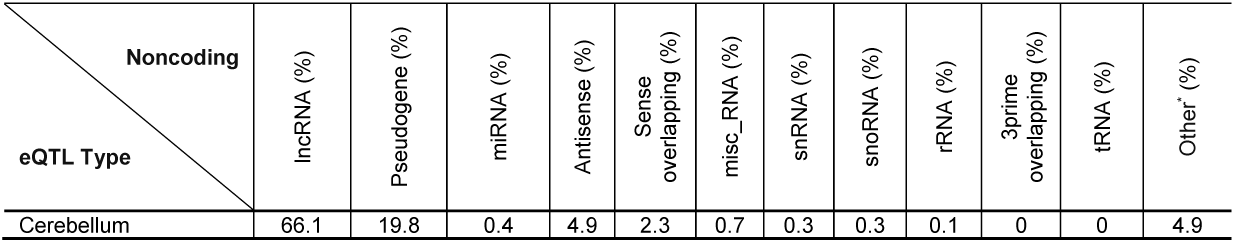
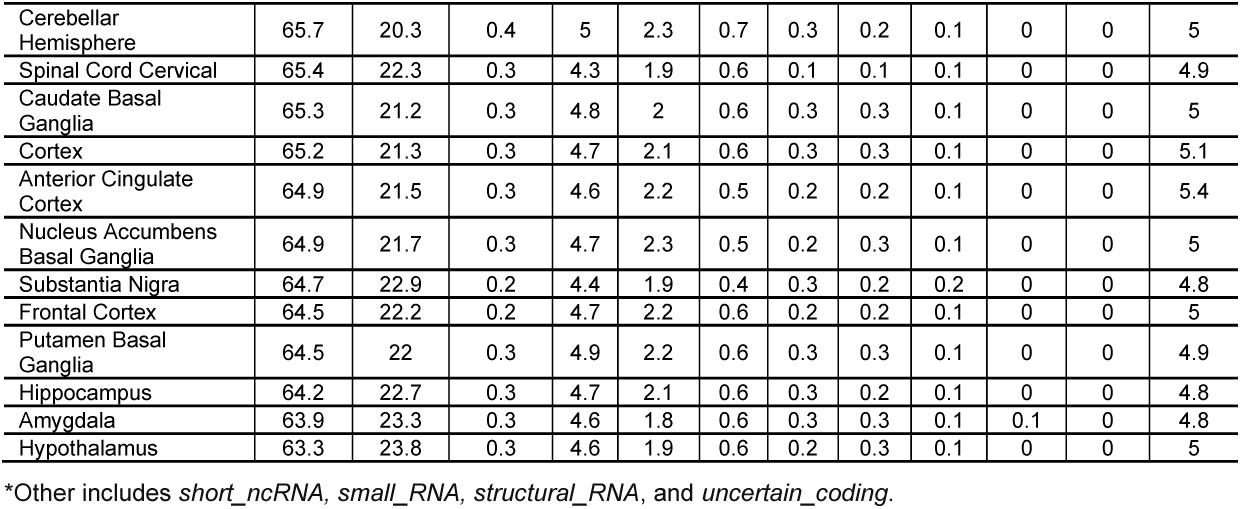
List of different noncoding RNAs fractions for every eQTL types (ncRNAs which contain at least one nervous system related eQTL polymorphism)

**Figure 3.**
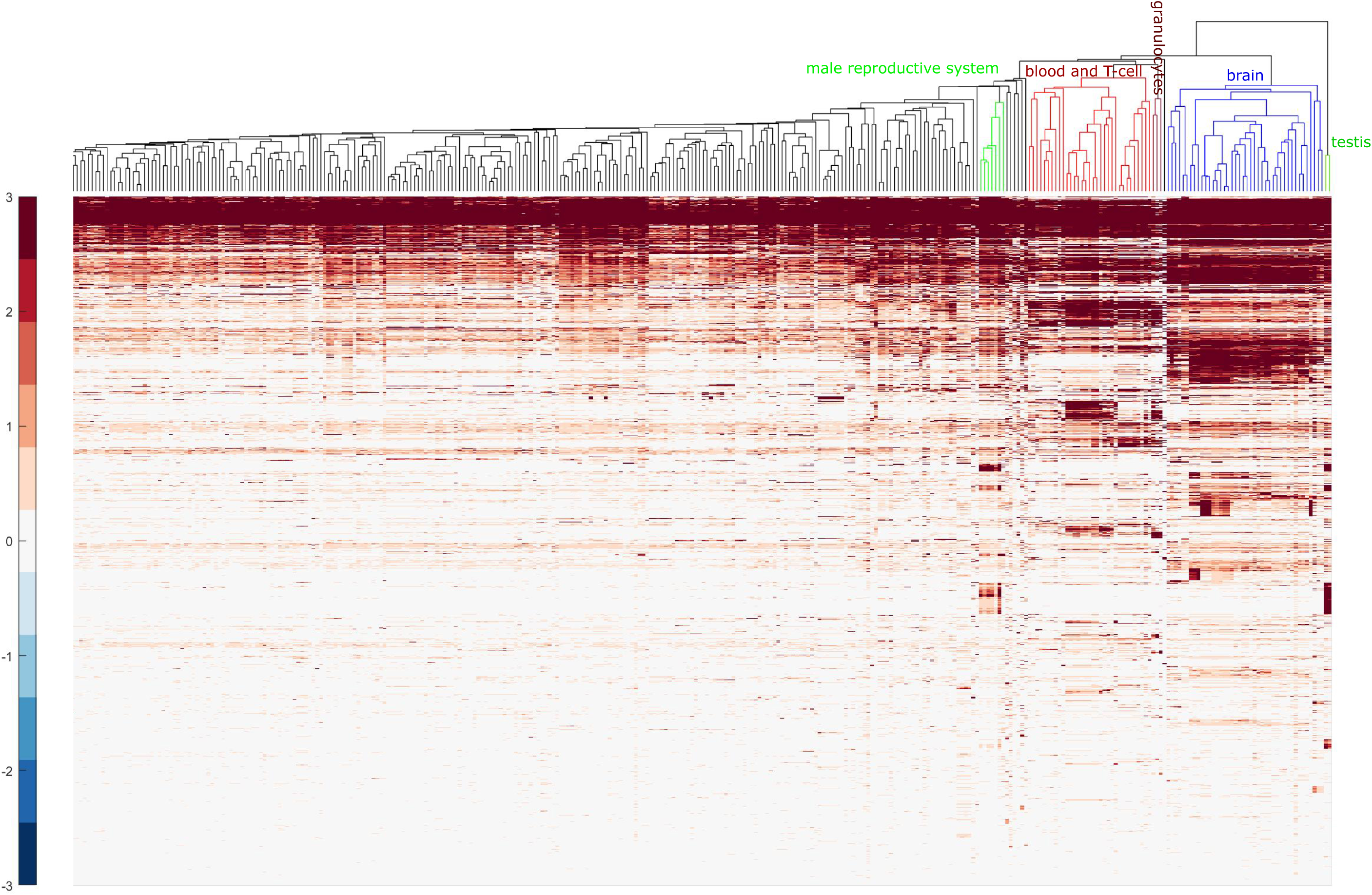
Expression profile of 17,743 noncoding RNA genes. Noncoding RNA genes that encompass at least one nervous system related eQTL polymorphism contain proportionally more nervous system enriched genes than expected. Hierarchical clustering of the 17,743 noncoding RNA genes based on their expression profiles revealed a substantial proportion of these genes were highly expressed in the brain while smaller clusters of genes were highly expressed in blood and T-cells tissues.

Of the 17,743 noncoding RNAs characterised with at least one nervous system-related eQTL polymorphism (**Supplementary table S2**), we find interactions with 0.47 coding genes per ncRNA (8,399 coding genes, with interactions defined as more than 0.002kb and less than 2,213kb from ncRNA, and with the average distance of 170kb - **Supplementary table S4**). These ncRNAs are interacting with an average of 0.08 noncoding genes (1,466 non-coding gene interactions identified).

Out of 17,743 noncoding RNAs (encompassing at least one nervous system related eQTL polymorphism), 14,530 ncRNAs interact with at least one protein-coding gene through eQTL pair including 1,866 in Amygdala, 3,718 in Anterior Cingulate Cortex, 5,258 in Caudate Basal Ganglia, 6,383 in Cerebellar Hemisphere, 8,675 eQTL pairs in Cerebellum, 5,869 eQTL pairs in Cerebral Cortex, 4,313 eQTL pairs in Frontal Cortex, 2,741 eQTL pairs in Hippocampus, 4,571 eQTL pairs in Hypothalamus, 4,571 eQTL pairs in Nucleus Accumbens Basal Ganglia, 3,711 eQTL pairs in Putamen Basal Ganglia, 2,229 eQTL pairs in Spinal Cord Cervical, and 1,410 eQTL pairs in Substantia Nigra (**Supplementary table S5**). The distribution of brain eQTL polymorphisms on different types of noncoding genes is illustrated in **Supplementary table S6**.

### Identify brain-enriched noncoding genes

We next used the FANTOM CAGE (Cap Analysis of Gene Expression) associated transcript database [16] and the FANTOM5 expression atlas [20] to characterise tissue-specificity features for noncoding genes. Of the 17,743 noncoding RNAs analysed, 7,164 are significantly expressed in nervous systems tissue (*see Methods*). Notably, the overwhelming majority (99.9%, 7163 out of 7164) are lncRNAs, with the exception of ENSG00000214857, which is a pseudogene (**Supplementary table S7**).

Next, we characterised brain-related noncoding RNAs with enhancer activity, based on the following criteria: these noncoding RNAs (i) are significantly enriched in brain tissue expression; (ii) are overlapping ENCODE-predicted chromatin states which identify enhancer and promoter loci; and (iii) are mapping to eQTL polymorphisms that interact with at least one protein-coding gene. Based on these criteria, we found 908 of 60,681 noncoding RNAs annotated as putative enhancers in brain tissue, which include 907 lncRNA and 1 pseudogene (**Supplementary table S8**). We refer to these as putative enhancer ncRNAs (enhancer-ncRNAs).

### Annotating ncRNAs with genome-wide association study SNPs

Next, we investigated these 908 putative “enhancer-ncRNAs” for potential regulatory function in neurodevelopmental disorders. To achieve this, we asked if enhancer-ncRNAs were associated with GWAS SNPs or CNV deletion or duplication loci documented in NDD patients. As a corollary, it has been reported that more than 96% of disease associated variants detected in genome-wide association studies (GWAS) map to the noncoding regulatory regions of genomes [21], suggesting a role in human disease. We found that of 908 enhancer-ncRNAs, 89 of these (9.8%) contain at least one GWAS SNP related to a neurodevelopmental disorder (**Supplementary table S9**). This is higher (P-value: 8.0e-02), when compared against 1,496 (8.4%) noncoding genes with at least one nervous system related eQTL polymorphism in our dataset (**Supplementary table S10**), and is also significantly higher (P-value: 1.9e-14) when compared to the 2,390 (4.8%) of all 60,681 noncoding genes which contain at least one GWAS SNP. **Table 2** shows the proportion of GWAS SNPs across neurodevelopmental disorders relevant to these 89 enhancer noncoding genes. Of these, we found that 68 lncRNAs are relevant to Schizophrenia-related GWAS SNPs, while 10 lncRNAs are associated with Psychosis, 9 lncRNAs with Attention Deficit Hyperactivity Disorder (ADHD), 9 lncRNAs Cognitive Impairment, 7 lncRNAs with Abnormality of the Nervous System, 2 lncRNAs Autistic Behavior, 2 lncRNAs Abnormality of emotion, 1 lncRNA Abnormality of metabolism, 1 lncRNA Abnormality of the endocrine system, and 1 lncRNA Language impairment. Notably, we find 20 lncRNAs which are associated with GWAS SNPs categorised to two or more neurodevelopmental disorders (**Table 2** and **Supplementary table S9**).

**Table 2.** List of neurodevelopmental disorders related GWAS that overlapped with the enhancer noncoding genes.

| Disease | Number of GWAS | % |
| --- | --- | --- |
| Abnormal emotion/affect behavior | 2 | 2.2 |
| Abnormality of metabolism | 1 | 1.1 |
| Abnormality of the endocrine system | 1 | 1.1 |
| Abnormality of the nervous system | 7 | 7.9 |
| Abnormality of the skin | 0 | 0.0 |
| Attention deficit hyperactivity disorder | 9 | 10.1 |
| Autistic behavior | 2 | 2.2 |
| Cerebellar hypoplasia | 0 | 0.0 |
| Cognitive impairment | 9 | 10.1 |
| EEG abnormality | 0 | 0.0 |
| Language impairment | 1 | 1.1 |
| Psychosis | 10 | 11.2 |
| Schizophrenia | 68 | 76.4 |
| More than two disorders | 20 | 22.5 |

Interestingly, we also found that for 56 enhancer-ncRNAs (out of 68 noncoding genes related to schizophrenia) there is no GWAS SNP in their eQTL linked coding gene. For non-schizophrenia NDDs (which we term “other NDDs”), we find 24 enhancer-ncRNAs in which there was no GWAS SNP detected in their eQTL linked coding gene (**Table 3**). Equally noteworthy is our finding that 14 ncRNAs for schizophrenia represent important associative loci, since they map to GWAS SNP loci relevant to neurodevelopmental disorder, while their eQTL genes do not contain a GWAS SNP signature, nor does their eQTL genes have interactions with any other eQTL data with a GWAS SNP (**Table 3**). These criteria also identify 8 ncRNAs as novel factors for NDDs which do not include schizophrenia (**Table 3**). These data suggest that noncoding RNA may be relevant to modulating expression in these genes, with relevance to neurodevelopmental disorders.

**Table 3:**
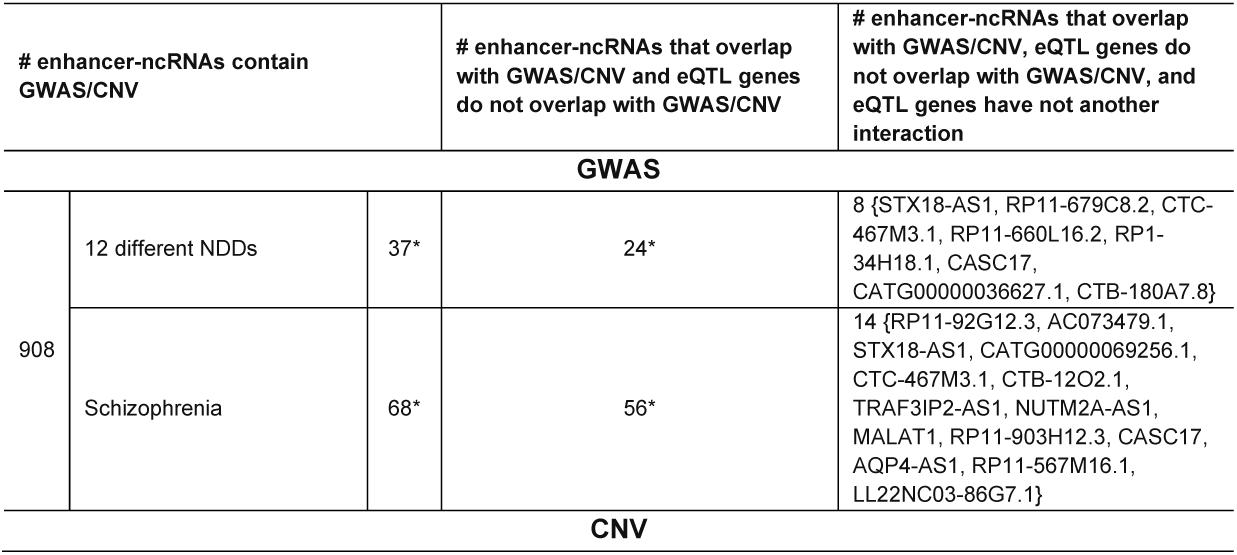
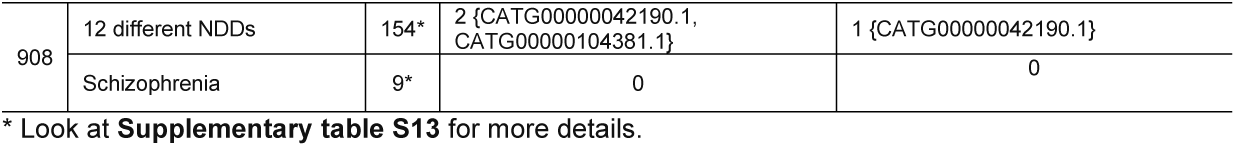
Summary of the GWAS/CNV overlapping analysis for enhancer ncRNAs.

### Investigating noncoding RNAs for an association with ASD or SCZ CNV loci

We investigated the overlap between 908 enhancer-ncRNAs (**Supplementary table S8**), and CNVs associated with Autism Spectrum Disorder (ASD) and Schizophrenia (SCZ). For ASD CNV associations, we utilised our recently reported tool SNATCNV [22] to identify high-confidence CNV loci in a cohort of 19,663 autistic and 6,479 non-autistic individuals (http://autism.mindspec.org/autdb; download date: July 2018), we mapped enhancer ncRNAs to loci for 3,260 deletion and 4,642 duplication regions, respectively (99% confidence, P-value threshold of 0.001; see *Methods section*). A full list of ASD associated CNVs identified by SNATCNV is provided in **Supplementary table S11**. For our investigation of enhancer-ncRNA associations with CNV loci for Schizophrenia, we performed SNATCNV analysis on a cohort of 21,210 SCZ individuals in which 262 deletion and 138 duplication loci were identified, respectively (**Supplementary table S12**). For each dataset, we identified enhancer-ncRNAs that overlap with ASD- or SCZ-associated CNVs, defined as loci with at least 10% positional overlap; *see Methods section*). Of the 908 enhancer ncRNAs analysed, we found 154 ncRNAs (17%) that were associated with ASD comprising 56 CNV deletion and 98 CNV duplication loci. In all, 9 ncRNAs (1%) are identified for SCZ in this investigation, comprising seven deleted and two duplicated CNVs (**Supplementary table S13**). We also performed the analysis on 17,743 noncoding genes (with at least one nervous system related eQTL polymorphism) (**Supplementary table S2**). We found that 2,572 noncoding genes (14.5%) overlap with at least one ASD-associated CNVs (including 936 deletions and 1,636 duplications). For schizophrenia, there were 167 ncRNAs (0.94%) including 122 associated with CNV deletions and 45 associated with CNV duplications types. When considering all 60,681 noncoding genes, we identified 8,787 (14.5%, including 2,577 deletions and 6,210 duplications) for ASD CNVs and 348 (0.57% including 269 deletions and 79 duplications) for schizophrenia. Thus, enhancer ncRNAs are more highly associated with CNV loci for ASD, compared to schizophrenia.

### Enhancer ncRNAs are associated with CNV loci for ASD and SCZ, and influence novel downstream genomic loci in different ways

From this analysis, we found that for two noncoding genes *CATG00000042190* and *CATG00000104381* (out of 154 noncoding genes related to ASD), their eQTL associated coding genes (*OR2W3* and *GDI2* linked with ncRNAs *CATG00000042190* and *CATG00000104381*, respectively) do not map to CNV loci. Furthermore, we found that only one ncRNA (*CATG00000042190*) maps to an ASD CNV whereas its eQTL-associated gene does not implicate an NDD CNV, and its eQTL gene (*OR2W3*) does not have another reported interaction through any eQTL from ASD CNVs (**Table 4**). This suggests that the enhancer-ncRNAs *CATG00000042190* and *CATG00000104381* are unique in their association with ASD, while their regulatory actions lie in downstream loci that have not previously been associated with this condition. In addition, for 268 of 908 enhancer ncRNAs, we find an association with NDD-related GWAS SNPs or ASD/SCZ associated CNVs (**Supplementary table S13**). Interestingly, for 82 of these, the eQTL associated coding gene does not overlap with either NDD related GWAS SNPs or ASD/SCZ associated CNVs (**Supplementary table S14**), suggesting the genomic variant in the enhancer-ncRNA may influence the expression of eQTL linked gene. Importantly, for 23 of 82 the eQTL linked coding gene does not overlap with GWAS/CNV, and also has no eQTL interaction with another coding or noncoding genes, including 14 enhancer-ncRNAs in SCZ and 9 enhancer-ncRNAs in other NDD associated genes (**Table 4**). These enhancer-ncRNAs are notable for their unique features. For example, *CTB-180A7.8* is a long noncoding RNA located on chr19:6393588-6411788. There are robust signals for HMM predicted enhancers, H3K27ac, H3K4me1 histone active marks, and DNase I hypersensitive sites overlapped with *CTB-180A7.8*. This brain-enriched lncRNA (brain-enriched expression P-value: 1.08e-73) is linked with gene *ALKBH7* through an eQTL link (**Figure 4**). Importantly, this lncRNA encompasses an ASD related GWAS SNP (rs6510896), and also lies within a region which is significantly deleted in ASD cases compared to controls (**Figure 4**). Of note, *ALKBH7* has not yet been associated with NDD related disorders, yet our findings suggest that enhancer-like activity of *CTB-180A7.8* for *ALKBH7*, as well as mutation or deletion in lncRNA *CTB-180A7.8* may be relevant to *ALKBH7* expression in neurodevelopment, and NDDs such as ASD.

**Table 4:** Final list of noncoding RNAs that overlapped with NDDs or schizophrenia related GWAS/CNVs.

| variant type | ncRNA chr | ncRNA start position | ncRNA end position | ncRNA symbol | eQTL linked genes |
| --- | --- | --- | --- | --- | --- |
| **NDDs** |  |  |  |  |  |
| CNV | chr1 | 247094914 | 247098641 | CATG00000042190 | OR2W3 |
| GWAS | chr4 | 4543868 | 4822163 | STX18-AS1 | EVC |
| GWAS | chr4 | 120987634 | 121338622 | RP11-679C8.2 | MAD2L1, C4orf3 |
| GWAS | chr5 | 87970840 | 88063654 | CTC-467M3.1 | TMEM161B |
| GWAS | chr11 | 71159674 | 71163869 | RP11-660L16.2 | KRTAP5-7, KRTAP5-8, KRTAP5-10 |
| GWAS | chr12 | 77717925 | 78222393 | RP1-34H18.1 | ZDHHC17 |
| GWAS | chr17 | 68608010 | 69739611 | CASC17 | SOX9 |
| GWAS | chr18 | 74696810 | 74700868 | CATG00000036627 | ZNF236 |
| GWAS | chr19 | 6393588 | 6411788 | CTB-180A7.8 | SLC25A41, SLC25A23, ALKBH7, CLPP |
| **Schizophrenia** |  |  |  |  |  |
| GWAS | chr1 | 200638614 | 200663382 | RP11-92G12.3 | DDX59, KIF14 |
| GWAS | chr2 | 6072923 | 6143655 | AC073479.1 | DKFZP761K2322(ensemble) |
| GWAS | chr4 | 4543868 | 4822163 | STX18-AS1 | EVC |
| GWAS | chr4 | 86068874 | 86305742 | CATG00000069256 | CDS1 |
| GWAS | chr5 | 87970840 | 88063654 | CTC-467M3.1 | TMEM161B |
| GWAS | chr5 | 151304308 | 152109235 | CTB-12O2.1 | GLRA1, G3BP1, NMUR2 |
| GWAS | chr6 | 111803916 | 111924133 | TRAF3IP2-AS1 | KIAA1919, REV3L |
| GWAS | chr10 | 88963526 | 89102347 | NUTM2A-AS1 | NUTM2A, NUTM2D, FAM35A |
| GWAS | chr11 | 65263885 | 65276602 | MALAT1 | AP000769.1 |
| GWAS | chr14 | 21153505 | 21268108 | RP11-903H12.3 | RNASE3, RNASE4, RNASE6 |
| GWAS | chr17 | 68608010 | 69739611 | CASC17 | SOX9 |
| GWAS | chr18 | 24235508 | 24771692 | AQP4-AS1 | KCTD1 |
| GWAS | chr18 | 77336391 | 77360135 | RP11-567M16.1 | CTDP1 |
| GWAS | chr22 | 22292599 | 22300188 | LL22NC03-86G7.1 | TOP3B |

**Figure 4.**
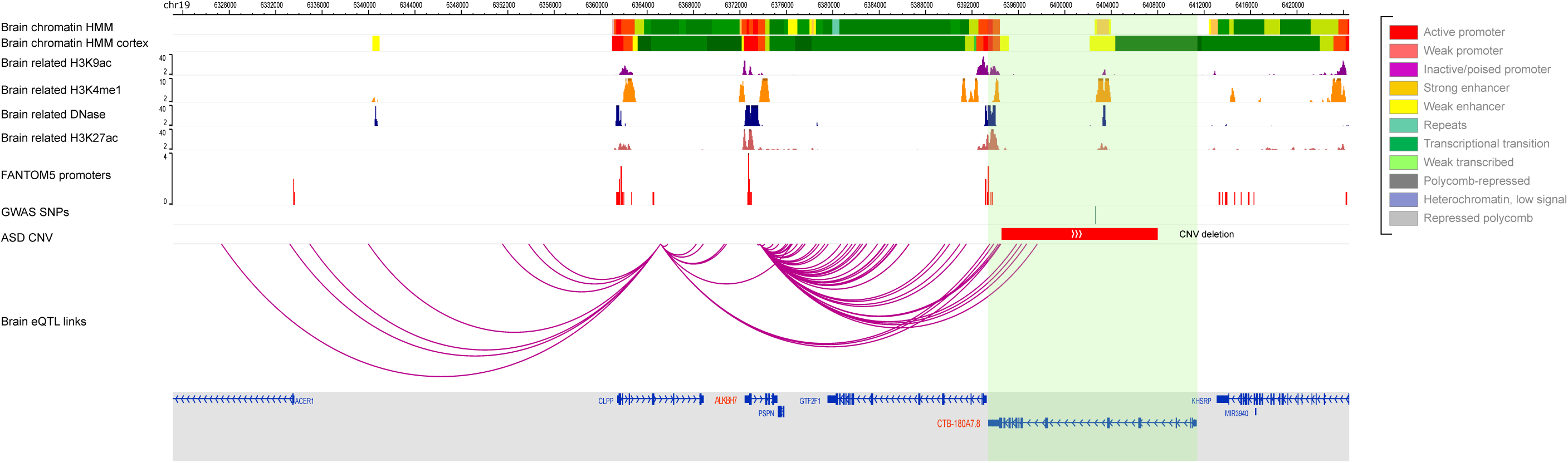
Another example loci of noncoding gene *CTB-180A7.8* interacting with eQTL linked gene *ALKBH7*. *CTB-180A7.8* overlaps with HMM predicted strong enhancer, strong signals of chromatin states H3K27ac, H3K4me1, and H3K9ac, all presented in brain. This lncRNA also overlaps with multiple FANTOM5 enhancers and contains one NDD related GWAS SNP. This lncRNA also significantly deleted in autistic individuals, while ***ALKBH7***, the eQTL linked gene, does not overlap with either GWAS SNPs or ASD related CNVs. All these observations may indicate a putative enhancer activity of this noncoding RNA.

In another example, the lncRNA *CATG00000053385* (chr20:45035426-45084326) interacts with the gene *SLC12A5* through an eQTL link. *SLC12A5* encodes potassium and chloride co-transporter 2 (KCC2), and is the main extruder of chloride in neurons that mediates the electrophysiological effects of GABA signalling [23]. Based on our observations, *CATG00000053385* may be relevant to modulation of *SLC12A5* expression, leading to NDDs, as follows. *CATG00000053385* overlaps with strong signals of HMM enhancers and histone active marks H3K27ac, H3K4me1, and DNase I hypersensitive sites, as detected in brain tissue analyses (**Figure 5**). Furthermore, this lncRNA is associated with two SCZ GWAS SNPs and one ASD related GWAS SNP (**Figure 5**), is significantly expressed in brain (brain-enriched expression P-value: 9.61e-09), and dynamically expressed in an induced pluripotent stem cell model of neuron differentiation. Collectively, these observations indicate that *CATG00000053385* is relevant to *SLC12A5* enhancer activity, and mutation in this lncRNA is associated with NDDs including ASD and schizophrenia.

**Figure 5.**
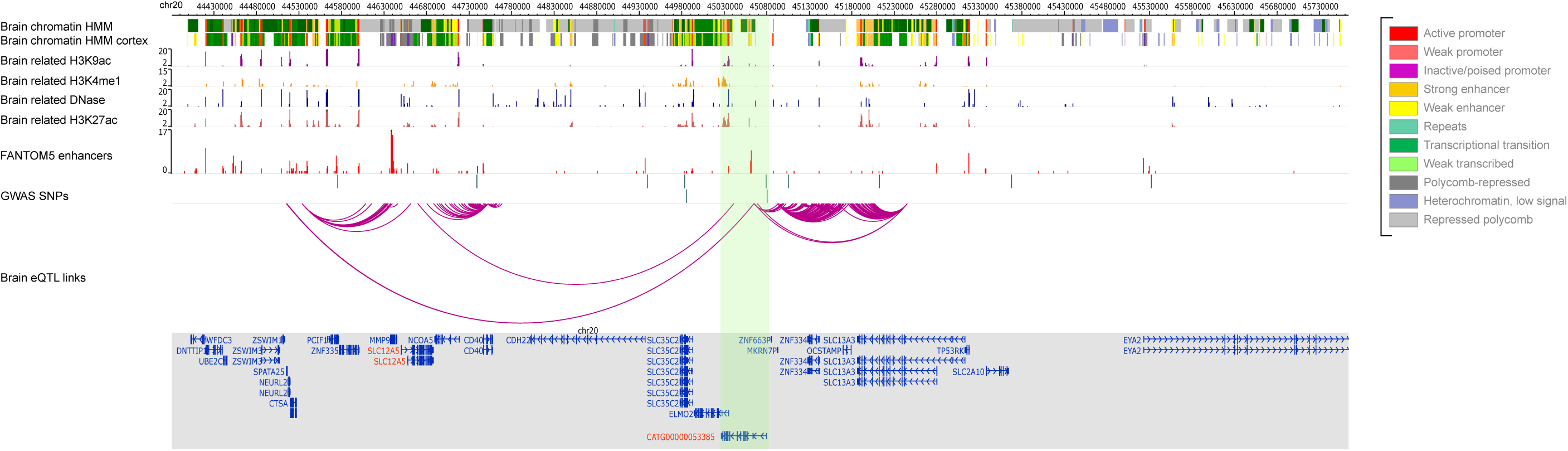
An example loci of noncoding gene *CATG00000053385* interacting with eQTL linked gene *SLC13A3*. *CATG00000053385* overlaps with HMM predicted strong enhancer, chromatin states H3K27ac, H3K4me1, and H3K9ac, all presented in brain. This lncRNA also overlaps with multiple FANTOM5 enhancers. All these observations may indicate a putative enhancer activity of this noncoding RNA. Interestingly, *CATG00000053385* encompasses two NDD related GWAS SNPs, while *SLC13A3*, the eQTL linked gene, does not overlap with either GWAS SNPs or ASD related CNVs. MHiC tool [72] has been used to generate the figure.

It is further noteworthy that 3 of 23 lncRNA genes are associated with related GWAS SNPs for both SCZ and NDDs, namely *STX18-AS1, CTC-467M3.1*, and *CASC17*. In the case of *STX18-AS1*, this lncRNA is located on human chromosome 4, and shows reduced expression in association with rs6824295, a SNP associated with heart disease [24], as well as rs16835979, whose clinical significance is unknown. Also, an eQTL polymorphism in *STX18-AS1* is associated with the coding gene *EVC*, yet we find no overlap of neither NDD related GWAS nor CNV associated with this coding gene. Interestingly, we identified two NDD related GWAS SNPs (rs2887447 and rs13130037) and one schizophrenia related GWAS SNP (rs7675915) overlapping with *STX18-AS1*, and also that *STX18-AS1* is significantly duplicated in individuals with Schizophrenia. Thus, mutations in the ncRNA *STX18-AS1* is relevant to *EVC* expression in NDDs. In the second example, *CTC-467M3.1* is a lncRNA located in chromosome 5, and is also known as *MEF2C* antisense RNA 2 (*MEF2C-AS2*). Although there is not any strong evidence for this gene to be involved in NDD or schizophrenia diseases, this gene is critical to neurodevelopment, learning and memory [25]. For example, *MEF2C* participates in a brain gene co-expression network which showed a promising therapeutic tool to persuade cognitive enhancement [26]– [28]. Finally, in the third example, the Cancer Susceptibility Candidate 17 (*CASC17*) is a non-protein-coding gene in chromosome 17 of the human genome. It was shown that genetic variants of *CASC17* gene could affect the risk for psychosis [29]. The *CASC17* gene also correlated with reactionary and impulsive behavior, given its characterisation as a high confidence candidate based on multiple SNPs identified through a genome-wide approach for children’s aggressive behavior [30].

### Association of enhancer ncRNAs with DECIPHER phenotypes

To better understand whether enhancer ncRNAs identified in this study are significantly associated with behavioural traits, we cross-correlated our results with patient phenotypes documented in DECIPHER (Database of Chromosomal Imbalance and Phenotype in Humans using Ensembl Resources) [31]. DECIPHER contains 1,094 patients annotated as autistic, 5,558 patients annotated with intellectual disability (552 both annotated as intellectual disability and NDD) and 6,513 patients with other phenotypes. To achieve this, we began by enumerating phenotypic terms in DECIPHER patients, with CNVs implicated in our 82 enhancer-ncRNAs (enhancer-RNAs that contain NDD related GWAS SNPs or CNVs but the eQTL linked gene does not contain NDD related GWAS SNPs or CNVs) which do not overlap with eQTL linked coding genes. The frequencies for pairs of enhancer-ncRNAs associated with a given DECIPHER phenotype were then calculated, to derive the proportion of DECIPHER patients with such criteria. To assess significance, we used 5,000 random permutations of the patient-phenotype associations in DECIPHER to estimate the fraction of patients with the same enhancer ncRNAs-phenotype pair that emerges by chance. Using this approach, we found 246 patients in DECIPHER where CNV overlapped with at least one of our enhancer ncRNAs, while the eQTL linked gene did not overlap with the CNV.

When we looked at this data further, we identified 48 enhancer ncRNAs (48 of 82 enhancer-ncRNAs), with at least one phenotype term over-represented (95% confidence interval in DECIPHER patients with CNVs overlapping the enhancer ncRNAs (**Supplementary table S15**). For 34 of enhancer-ncRNAs we identified at least three DECIPHER patients with CNVs overlapping the enhancer ncRNAs (**Supplementary table S17**). For example, enhancer ncRNA *ERICH1-AS1* is interacting with coding gene *FBXO25*, which has been associated to psychiatric phenotypes (PMID: 31849056). We found both NDD and SCZ related GWAS mutations in *ERICH1-AS1* but not in *FBXO25*. Our DECIPHER analysis revealed 13 samples in the DECIPHER database with CNVs that encompassed our candidate ncRNAs *ERICH1-AS1* and did not overlap with eQTL linked coding genes *FBXO25*. Interestingly, phenotypic terms Autism, Intellectual Disability, and Macrocephaly were enriched for DECIPHER patients with CNV overlapping with *ERICH1-AS1* (**Supplementary table S17**). Another example is enhancer-ncRNA *SYN2*. Nervous system related eQTL data shows that this ncRNA interacts with protein-coding gene *PPARG*. Our literature search showed that still there is no a consensus about the function of *PPARG* in NDD/SCZ disorders (PMID: 29229843, PMID: 19560328). However, we found *SYN2* as a potential enhancer for *PPARG* that contains NDD related GWAS mutation. Interestingly, we found seven samples in the DECIPHER database with CNVs that encompassed our candidate ncRNAs *SYN2* and did not overlap with eQTL linked coding genes *PPARG*. The DECIPHER analysis also revealed that HPO terms “Delayed Speech and Language Development”, “Seizures”, and “Abnormality of the Ear” were enriched within these seven DECIPHER patients. This would suggest that genomic variant in *SYN2* may be responsible for the functionality of *PPARG* in NDD/SCZ disorders. Thus, for these enhancer-RNAs, we find unique associations with known neurodevelopmental genes, as well as previously uncharacterised candidates.

### Investigating the impact of 82 enhancer-ncRNAs for functions in neurodevelopment and NDDs

Our investigation has led to the identification of 82 enhancer ncRNAs with an association with NDD related GWAS SNP or ASD/SCZ CNV, and which have eQTL-associated coding genes that do not overlap with neither NDD-related GWAS SNPs nor ASD/SCZ CNV loci (**Supplementary table S14**). When we conducted a literature search for evidence to support the hypothesis that these enhancer-ncRNAs influence gene expression relevant to nervous system development and NDDs, we identified four notable examples, as follows.

In the first example, we identify *MALAT1* (Metastasis Associated Lung Adenocarcinoma Transcript 1) as an NDD-related enhancer-ncRNA, located within chromosome 11 with 12.7kb in length; this lncRNA is highly expressed in neurons [32]. It has been shown that the loss of the *MALAT1* negatively affects the expression of neighboring genes, including genes which are significantly decreased in expression within several brain regions relevant to schizophrenia [33]. Furthermore, a study reported that *MALAT1* expression was significantly different in patients with schizophrenia compared with healthy subjects [34], and *MALAT1* is also implicated in ASD [35]. In addition, the brain eQTL data shows *MALAT1* has an interaction with a protein-coding gene (*AP000769.1*), although *AP000769.1* is associated with bone mineral density [36], while its function with NDDs is uncharacterised. In the second example, we find that *SYN2* (Synapsin II) is a candidate gene for NDD which is influenced by enhancer-ncRNAs. This gene is 253.9kb in length and located in chromosome 3, and is a member of the synapsin gene family. One of its variants belongs to a noncoding gene class. *SYN2* is associated with abnormality of the nervous system through the GWAS data [37], and its association with schizophrenia is reported in several studies, particularly in families of Northern European ancestry [38]–[40]. Studies also indicate that *SYN2* loss-of-function may be associated with autism, abnormal behavior, and epilepsy in humans, and in comparable neuropathological features observed in mice [37]. *SYN2* interacts with four different protein-coding genes (*PPARG, TIMP4, RAF1, and CAND2*), as indicated in the brain eQTL data. Yet, for *PPARG*, its association with schizophrenia remains to be clarified, owing to conflicting results [41], [42]. Also, there is evidence that *PPARG* is associated with early postnatal neurodevelopmental disease and cerebral connectivity [43], [44]. In the case of *TIMP4*, this gene is associated with both ASD and schizophrenia [45]. Similarly, *RAF1* and *CAND2* are implicated in NDD and schizophrenia [46]–[49].

The third example of an enhancer-ncRNA with unique features is *SOX2-OT* (SOX2 overlapping transcript (*SOX2OT*)). This lncRNA is 847.1kb in length, located in chromosome 3 and comprises at least 5 exons. Our analysis showed that there are nine GWAS SNPs associated schizophrenia within *SOX2-OT*. It is an important regulator of neurogenesis and is dynamically expressed during neural cell differentiation [50], and *SOX2OT* is associated with schizophrenia, as reported in a study of 108 schizophrenia-associated genetic loci [51], [52]. In addition, it has been shown that *SOX2OT* is relevant to ASD as well [53], [54]. Based on the brain eQTL data, *SOX2OT* interacts with two protein-coding genes (*CCDC39* and *FXR1*), of which *CCDC39* is associated with schizophrenia [52], while *FXR1* may be relevant in mental conditions [55], [56].

The fourth example of an enhancer-ncRNA for NDD is *ATXN8OS* (Ataxin 8 opposite strand (*ATXN8OS*)), a lncRNA gene 23.7kb in length, and located in chromosome 13. This gene is associated with cognitive impairment in a GWAS SNP study [57], and is implicated in schizophrenia through a study with the GeneAnalytics program [58]. Furthermore, *ATXN8OS* is suggested to be relevant to neurodegenerative diseases through dysregulation of gene expression [59], [60]. *ATXN8OS* interacts with a protein-coding gene *KLHL1* based on the brain eQTL data, and *KLHL1* is associated with gait disturbance, a motor phenotype affecting some individuals with ASD [61]. Furthermore, *KLHL1* is reported to have a co-expressed interaction with *TSHZ3* gene which itself is associated with ASD [62]. Also, *KLHL1* is implicated in Asperger Syndrome, based on a GWAS study [63]. Taken together, we find that enhancer-ncRNAs influence a wide range of known and previously uncharacterised genes to modulate neurodevelopment and NDDs.

## Discussion

We developed an integrative bioinformatics pipeline to systematically analyse and identify noncoding RNAs with possible enhancer role in NDDs. Our pipeline analysed a large set of ncRNAs through a workflow which integrated ENCODE regulatory features, FANTOM5 expression profiles, and nervous system related eQTLs pairs to identify 908 ncRNAs as putative enhancers in NDDs. The 908 ncRNAs were then further evaluated with neurodevelopmental associated GWAS SNPs and Copy Number Variations to identify those putative ncRNA enhancers that significantly deleted/mutated or duplicated in neurodevelopmental samples against healthy individuals. Through this approach, we identified 82 putative ncRNA enhancers that contained NDD related genomic variants (GWAS SNP or CNV) and interact with eQTL-linked genes, but that no genomic variants overlapping with eQTL-linked genes. We called these putative ‘enhancer ncRNAs’. Interestingly, of these enhancer ncRNAs, for 23 of them, the noncoding-RNA is the only region interacting with the eQTL-linked gene. It suggests that deletion or duplication in these ncRNAs may affect the expression of the eQTL linked gene. Of 82 enhancer ncRNAs, our systematic literature search revealed that 18 ncRNAs are associated with neurodevelopmental disorders, while 15 are associated with schizophrenia. Furthermore, expression of 39 of the 82 enhancer ncRNAs are stimulated in an induced pluripotent stem cell model of neuron differentiation, suggesting their relevance to neurodevelopment. Our literature search further revealed that 28 of the 144 eQTL linked coding genes (after duplications removed) interacting with the 82 enhancer ncRNAs are NDD associated genes (**Supplementary table S16**). Hence, of these, we report the remaining 116 coding genes as novel NDD candidate genes which are associated with enhancer-ncRNAs.

One interpretation of our findings is that genomic variation in a noncoding RNA or its eQTL-mapped enhancer loci influences expression of downstream target genes enriched in brain tissues, leading to NDD. Of note, we find that enhancer-ncRNAs, and their corresponding eQTL are brain-enriched. Also, eQTL-linked coding genes (74 of 163 eQTL coding genes interacting with the ncRNAs) are also brain-enriched. Co-expression of these genes with ncRNAs harboring NDD-associated mutations may suggest that the noncoding RNAs are involved in similar biological functions and are potential candidate NDD risk loci. For example, the ncRNA *CATG00000070525* (chr4:149,363,864-149,495,900) interacts with brain-enriched coding gene *NR3C2*. The *NR3C2* is a member of a family of nuclear receptors in which their ligands diffuse into cells, interacts with their cognate receptor to signal gene expression changes within the nucleus. This brain-enriched gene is classified by SFARI gene as category 3 ‘Suggestive Evidence’ ASD candidate genes. As a corollary, mutation of the mouse ortholog *nr3c2* leads to nervous system phenotypes [64]. Also, our analysis of DECIPHER data found one DECIPHER sample with CNV deletion that overlaps with *CATG00000070525* did not overlap with eQTL-linked coding gene *NR3C2*. Thus, our investigation has identified a role for *CATG00000070525* to influence *NR3C2* in NDD.

By profiling post mortem brains (from 48 ASD, 49 controls) Parikshak *et al*. [65] identified 60 lincRNAs based on dysregulated expression, and not mutation in ASD patients, which may play a role in ASD, but are not causal genes *per se*. We analysed these data to find that only one of the reported lincRNAs from this study, the lincRNA ENSG00000271913, lies within an ASD CNV region, or contains an ASD-related GWAS SNPs. Yet, the relevance of ENSG00000271913 in ASD is confounded by the finding that it is recurrently co-deleted with *SHANK3*, a well-characterised ASD causal gene [66]. Of note, our brain-enriched enhancer-ncRNAs are significantly detected in NDD individuals with deletion/duplication or GWAS SNPs. it suggests disruption of interaction between these ncRNAs with eQTL-linked genes are most likely to be candidate for the observed phenotype.

Finally we interrogated the DECIPHER database [31] to identify patients with genomic abnormalities in the enhancer ncRNAs but not in the eQTL-linked genes. For 79 of 161 interactions (eQTL interaction enhancer-ncRNAs and eQTL coding genes), we identified candidates in which there was at least one DECIPHER sample in which a genomic abnormality is observed in the enhancer-ncRNAs, but not in the eQTL-linked genes. This led to the identification of 34 interactions, of which there were at least 3 patients for each (**Supplementary table S17**). Consistent with a role for our set of enhancer-ncRNAs in NDD, this analysis led us to identify overrepresented traits including autistic behavior, delayed speech and language, cognitive impairment and global developmental delay for 77 of the 79 DECIPHER samples. Our data therefore suggests that genomic variants in these enhancer ncRNAs are relevant to neurodevelopmental disorders, unique to their functions as modulators of enhancer activity. For 34 of the 82 enhancer ncRNAs, at least one phenotype term was over-represented (fold enrichment above 2) in DECIPHER patients. Of note, the majority of these 34 noncoding RNAs had unique combinations of overrepresented phenotypes (73 combinations in total). Therefore, genomic abnormalities for enhancer ncRNAs are associated with distinct clinical phenotypes. The categorisation of these combinations of phenotypes for enhancer-ncRNAs in this study may be informative in the phenotypic characterisation of patients (e.g. obesity, microcephaly, macrocephaly, split hand). Importantly, for patients with genomic abnormalities in enhancer–ncRNAs we have identified in this study, the associated co-morbidities that we identify, such as renal cysts, atrial septal defects and tetralogy of Fallot, may be relevant for clinical consideration in NDD presentation.

## Methods and Materials

### CNV data

For the CNV region discovery, autism CNV data from 19,663 autistic patients (47,189 CNVs) and 6,479 normal individuals (24,888 CNVs) compiled from multiple large NDD CNV datasets by the Simons Foundation Autism Research Initiative (SFARI) was downloaded (http://autism.mindspec.org/autdb; download date: July 2016). Schizophrenia CNV data from 27,034 schizophrenia and 25,448 control individuals where obtained from the Marshall *et al*. study [67] (downloaded from the European Genome-Phenome Archive with accession number EGAS00001001960).

### Gene list

We have combined two genes list from FANTOM5 [16] and Ensemble [68] consortia (genome building hg19). This enables us to have a comprehensive list of noncoding genes. An in-house script has been used to combine the lists based on gene coordinates and/or gene names. The combined list can be found in the **Supplementary Table S18**). In total, 84,065 genes are available in the final list including 60,681 noncoding genes. **Figure 2** shows distribution of different types of noncoding genes in the list.

### Identification of genes with nervous system enriched expression

We used the list of brain-enriched genes in our previous paper [22]. In summary, to determine whether a gene had enriched expression in the 101 nervous system samples profiled by the FANTOM5 consortium (compared to the total set of 1,829 samples profiled by FANTOM), we first ranked samples by expression and then selected the top 101 samples most highly expressing the gene. We then used Fisher’s exact test to determine a P-value indicating whether the top 101 samples were more likely to be nervous system samples or not. To determine a threshold on the P-value we carried out 50,000 permutations randomising the sample labels to determine P-value thresholds at the 99% confidence interval; any gene with a P-value better than this permutation-specific P-value threshold, we consider a significantly nervous system-enriched gene.

### ChromHMM predicted promoter and enhancer regions

We have downloaded ChromHMM predicted promoters and enhancers from the ENCODE project [69]. First, we have used an in-house script in order to explore ChromHMM results for each gene. However, there are eight different ChromHMM dataset from human brain tissue samples. So, we used the consensus term of the eight ChromHMM that overlapped with a gene. For each region, ChromHMM data could be the following terms: active TSS, flanking active TSS, transcr at gene 5’ and 3’, strong transcription, weak transcription, genic enhancers, enhancers, ZNF genes repeats, heterochromatin, bivalent poised TSS, flanking bivalent TSS, bivalent enhancer, repressed polycomb, weak repressed polycomb, and quiescent low. We have used the maximum repeated term (from eight different samples), as a consensus term for each gene. If more than one term is shown, all of the equivalent terms have been considered.

### Overlapping between noncoding RNAs and eQTL pairs

eQTL data were downloaded from the Genotype-Tissue Expression (GTEx) Project [19] for thirteen different tissues of brain (Amygdala, Anterior Cingulate Cortex, Caudate Basal Ganglia, Cerebellar Hemisphere, Cerebellum, Cortex, Frontal Cortex, Hippocampus, Hypothalamus, Nucleus Accumbens Basal Ganglia, Putamen Basal Ganglia, Spinal Cord Cervical, Substantia Nigra). The total number of entries for each location is shown in **Supplementary Table S19**. We used an in-house script to find noncoding genes that encompass eQTL polymorphisms.

### GWAS data

GWAS SNPs were downloaded from GWAS Catalog (https://www.ebi.ac.uk/gwas/docs/file-downloads) and GWASdb v2 (http://jjwanglab.org/gwasdb). We only considered those SNPs that were associated with phenotypes Abnormal Emotion/Affect Behavior, Abnormality of Metabolism, Abnormality of the Endocrine System, Abnormality of the Nervous System, Abnormality of the Skin, Attention Deficit Hyperactivity Disorder, Autistic Behavior, Cerebellar Hypoplasia, Cognitive Impairment, EEG Abnormality, Language Impairment, Psychosis, and Schizophrenia.

### Annotate noncoding RNAs with GWAS using PLINK

As GWAS SNPs were reported using different genome build versions hg19 and hg38, we first converted all GWAS SNP coordinates to UCSC hg19 using UCSC Lift Genome Annotations tools [70] and confirmed the locations with NCBI remap tools (www.ncbi.nlm.nih.gov/genome/tools/remap). An in-house script has then been used to combine both datasets based on gene coordinates and/or gene names. The combined GWAS list can be found in **Supplementary table S20**. In order to perform a GWAS annotation analysis using PLINK [71] it is needed to prepare three files; an association file (based on our GWAS list), an attribution file downloaded from dbSNP (build 129), and a gene list file (created based on our gene list). We have also considered ±1Kbp distance from a gene as well.

### Using SNATCNV to identify CNVs associated with NDD

We used SNATCNV to identify significantly deleted or duplicated CNVs in NDD and SCZ cohorts. In summary, To identify regions that are significantly more often deleted or duplicated in autism patients than in phenotypically normal individuals, SNATCNV counts, for every position in the genome, how often the base was either deleted or duplicated in the case and control CNVs. Fisher’s exact test is then used to determine whether the base was significantly more frequently observed in a duplicated or deleted region of cases or controls. To determine the significance of this association SNATCNV used permutation testing by randomising 500,000 permutations for a range of observed CNV frequencies. In this study, we defined ‘significant regions’ as those with P-values better than 0.001, corresponding to 99% confidence. We next performed the same analysis to identify significantly deleted or duplicated CNVs in SCZ cohort. We then considered genes as deleted or duplicated if they had at least 10% overlap with the significant NDD associated CNVs.

### Annotation of known and putative novel NDD associated genes

For the candidate ncRNAs annotated as putative enhancer identified in this study, we carried out a comprehensive literature search of all genes that are interacting with the noncoding RNAs. For the extended literature search, we specifically searched for “gene name + autism” or “gene name + NDD” (e.g. *SAMD11* + autism or *SAMD11* + NDD), or “gene name + enhancer”. As a result, if a gene had a previous enhancer casual association, it was labelled as ‘there is a known enhancer’.

## Author contributions

HAR designed the study; HAR, YA, and JI-TH wrote the manuscript; HAR, JI-TH, YA, ARRF, and NL edited the manuscript. YA and HAR carried out all the analyses including the statistical analyses, region identification, text mining, eQTL pair analysis, GWAS, DECIPHER, and gene ontology. HAR and YA curated the genes and regions. YA created figures 1, 2 and all tables except tables S11, S12, S15, S16, S17, S19, S20. HAR generated figures 3, 4, and 5 and tables S11, S12, S15, S16, S17, and S19. All authors have read and approved the final version of the paper.

## Ethics approval and consent to participate

NA

## Availability of data and materials

All data are publicly available and cited in the paper.

## Consent for publication

All authors have read and approved the final version of the paper.

## Declaration

Declare in any published work that those who carried out the original analysis and collection of the Data bear no responsibility for the further analysis or interpretation of the data.

## Acknowledgments

We kindly acknowledge Government of Western Australia, Department of Health, Clinical Excellence, for their support of this project through MERIT award to HAR. This study makes use of data generated by the DECIPHER community. A full list of centres who contributed to the generation of the data is available from https://decipher.sanger.ac.uk/about/stats and via email from. Funding for the DECIPHER project was provided by Wellcome.

## Supplementary tables legends

**Supplementary table S1:** Coding and noncoding RNAs that overlapped with histone mark and ChromHMM signals.

**Supplementary table S2:** Results of overlapping noncoding RNAs with brain-related eQTL polymorphism.

**Supplementary table S3:** Statistics related to the brain eQTL genes that interact with coding and noncoding genes.

**Supplementary table S4:** Statistics of minimum, maximum, and average distance of eQTL interactions for noncoding RNAs with at least one nervous systems related eQTL polymorphism.

**Supplementary table S5:** List of noncoding RNAs with at least one nervous systems related eQTL polymorphism.

**Supplementary table S6:** Distribution of brain eQTL polymorphisms on different types of noncoding genes.

**Supplementary table S7:** List of brain-enriched noncoding genes.

**Supplementary table S8**: List of noncoding RNAs as putative enhancers based on the criteria used in the study.

**Supplementary table S9:** List of putative enhancer ncRNAs that contains neurodevelopmental related GWAS SNP.

**Supplementary table S10:** Results of overlapping noncoding genes with at least one nervous system related eQTL polymorphism with neurodevelopmental related GWAS SNP.

**Supplementary table S11:** A full list of neurodevelopmental related CNVs identified by SNATCNV.

**Supplementary table S12:** A full list of schizophrenia related CNVs identified by SNATCNV.

**Supplementary table S13:** List of putative enhancer ncRNAs that overlap with neurodevelopmental- or schizophrenia-associated CNVs.

**Supplementary table S14:** Literature search for enhancer-ncRNAs.

**Supplementary table S15:** Decipher analysis.

**Supplementary table S16:** Literature search for 144 eQTL linked coding genes interacting with the 82 enhancer ncRNAs.

**Supplementary table S17:** Decipher analysis for enhancer-ncRNAs with at least three evidences in DECIPHER.

**Supplementary table S18:** The combined list of genes used in this study.

**Supplementary table S19:** Total number of entries for eQTL data of thirteen different tissues of brain.

**Supplementary table S20:** The combined list of GWAS used in this study.

